# Transcriptomic profiling of desert tree *Prosopis cineraria* under heat stress reveals potential role of multiple gene families in its high thermotolerance

**DOI:** 10.1101/2025.05.31.652403

**Authors:** Hansa Sehgal, Rita A. Sharma, Mukul Joshi

## Abstract

The static nature of plants restrains their potential to evade heat stress and requires them to withstand stress through inherent defence abilities. *Prosopis cineraria* is a leguminous phreatophyte distributed across arid and semi-arid regions of India and can tolerate very high temperatures due to its adaptive physiological and biochemical mechanisms. Therefore, *P. cineraria* represents a repository of genes for abiotic stress tolerance. Two-months-old *P. cineraria* plants were subjected to heat stress at two different temperature regimes and transcriptome sequencing was performed to identify differentially expressed genes (DEGs). A total of 1151 and 1562 DEGs were observed in response to 45℃ and 55℃ heat stress compared to control, respectively, indicating that 55℃ treatment has a pronounced effect on *P. cineraria*. The transcriptomic data highlighted the potential role of multiple gene families and their interactions for high thermotolerance of *P. cineraria.* The expression of a few representative heat stress-responsive genes was validated with real-time qPCR. The in-depth bioinformatic analysis provided the detailed transcriptome profiling, supported by its validation, and new insights for important abiotic stress-related genes from thermotolerant *P. cineraria,* which can be used for crop improvement.

## 1. Introduction

In the current scenario, global warming will continue to increase (2021-2040) due to increased CO_2_ emissions and the associated factors (IPCC, 2023). As a result, the mean annual temperature will rise by 1.5 degrees globally. Plants rely on an optimum range of temperature for growth and development (Mondal et al., 2023). Rise in temperature impacts the crop productivity and would significantly threaten food security. Heat stress (HS) can affect plants at any time during their development and negatively impact their growth. In comparison to other stages of growth, reproductive stage is highly sensitive to HS (Jagadish et al., 2021; Resentini et al., 2023). For agricultural productivity, it is important to understand how plants cope with heat stress. Plants, being sessile organisms, can’t escape or avoid stress conditions; nonetheless, they have evolved complex molecular networks allowing them to sense HS and modify their cellular pathway to adapt according to the environment (Haider et al., 2021; Sehgal et al., 2025). HS negatively affects the photosynthetic efficiency, respiratory enzyme activities, disrupts the movement of biomolecules, and alters the permeability and fluidity of the plasma membrane. In addition, HS affects the assimilation of carbon, the ATP-generating system, and chloroplast biosynthesis, leading to plant senescence (Fortunato et al., 2023; Kan et al., 2023). At the cellular level, HS results in a burst of reactive oxygen species (ROS), such as hydrogen peroxide. To control the levels of ROS production, the production of ROS-scavenging enzymes including peroxidase (POD), superoxide dismutase (SOD) and catalase (CAT); and non-enzymatic molecules including mannitol, ascorbic acid and vitamin E increases (Jagadish et al., 2021; Qian et al., 2022). Heat tolerance in plants involves crosstalk between different pathways and majorly involves expression of heat shock transcription factor (HSF) and heat shock proteins (HSPs) (Yang et al., 2024). HSPs are molecular chaperones classified according to molecular size into five categories; HSP100, HSP70, HSP60, HSP40 and HSP20. Previous studies have shown that HS transduces the expression of specific HSPs (Lee et al., 2022; Tan et al., 2024). Among HSPs, HSP20 is the most responsive to heat stress and accumulates immediately. In several plant species, transcriptomic studies have reported the presence of HSP20 under HS conditions (Wang et al., 2017; Wang et al., 2021). HS triggers mis-folding and denatured protein accumulation, causing the unfolded protein response (UPR) in the lumen of endoplasmic reticulum, thereby maintaining protein homeostasis. In plants, UPR mitigates heat damage via two signalling pathways that involve: 1) basic-leucine zipper transcription factors, and 2) RNA splicing factors and their target RNA. Apart from the HSF-HSP mechanism, plant’s endogenous phytohormones like abscisic acid (ABA), jasmonic acid (JA), ethylene, and salicylic acid (SA) play a crucial role in improving the tolerance of plants to HS (Ahammed, 2016).

*Prosopis cineraria*, a state tree of Rajasthan, India and also the national tree of the United Arab Emirates, holds paramount importance as a multifaceted tree species. *P. cineraria* belongs to the Fabaceae family, and is known to thrive under harsh conditions through unique genetic mechanisms, serving as an excellent source of genetic elements that confer tolerance under abiotic stress conditions (Sasi et al., 2024; Sharma & Singh, 2025). The genome of *P. cineraria* has been sequenced recently (Sudalaimuthuasari et al., 2022). However, study related to the response of *P. cineraria* against abiotic stresses and the pool of responsive genes is limited. So far only one gene, *PcHSP17.9*, has been characterised for its role during abiotic stress (Kaur et al., 2018). In recent years, the cost of genome sequencing has been significantly reduced. Therefore, RNA-sequencing technology has been extensively used to elucidate the response of the model and non-model plants towards abiotic stress and for the identification of unique or conserved stress-responsive genes (Rai et al., 2017). Transcriptomic studies provide key insights into the function and structure of genes. In this study, we have utilised RNA-seq approach to understand the genome-wide gene expression patterns and unravel the novel and conserved transcripts, transcription factors, and molecular response of *P. cineraria* plants to extreme heat stress. Characterization of *P. cineraria* heat responsive genes can pave the way to enhance the abiotic stress tolerance of non-thermotolerant plants.

## 2. Materials and methods

### 2.1 Plant growth, stress treatment and sample collection

*P. cineraria* seeds were collected from Central Arid Zone Research Institute, Jodhpur, India. Seeds were germinated in pots in controlled conditions of a greenhouse, and two-month-old plants of identical growth were taken for HS treatments. While 100 plantlets were kept under control conditions, a set of 100 plantlets each were subjected to heat stress treatment at 45°C and 55°C, respectively, for two hours per day (d) up to 10 d. Leaves of similar developmental stage were collected on 4^th^, 7^th,^ and 10^th^ d for biochemical analysis and molecular studies. Phenotypically, it was observed that *P. cineraria* plants could sustain 45°C temperature; their leaves were green, and the plants were healthy and comparable with control plants. These experiments suggest that *P. cineraria* is highly tolerant to HS and can easily live up to 45°C during its initial growth period.

### 2.2 RNA extraction, library preparation, and sequencing

*P. cineraria* leaves contain heavy mucilaginous content. Total RNA isolation was tried by using various methods, including the Trizol method, the GITC method, and the CTAB method, but RNA isolation was only partially successful with the CTAB method. The CTAB method was modified further to optimize good-quality RNA isolation. The total RNA was extracted from control and treated *P. cineraria* leaf samples (100 mg) using the modified CTAB method. This method included overnight incubation after lithium chloride treatment and an additional chloroform: isoamyl treatment step. RNA was collected from 7-d heat stress-treated samples, i.e., 25°C (Control), 45°C (T-45), and 55°C (T-55), with two biological replicates of each condition. The total RNA quality and quantity were checked with a NanoDrop ND-1000 Spectrophotometer and further evaluated on a 1.5% MOPS-Agarose gel. Before proceeding with library preparation, the integrity of the RNA was assessed on the Bioanalyzer. RNA samples with RNA integrity numbers ≥7 were considered high-quality and used further for library preparation and sequencing. Total 500ng of total RNA was used to enrich the mRNA using NEBNext^®^ Poly(A) mRNA magnetic isolation module (Catalog: E7490, New England Biolabs) by following the manufacturer’s protocol. The enriched mRNAs were further taken for library preparation using the NEBNext^®^ Ultra^™^ II RNA Library Prep Kit for Illumina (Catalog: E7775S, New England Biolabs). A total of six libraries were sequenced using the Illumina HiSeq 2X (150) bp Chemistry platform. Raw reads generated from sequencing were submitted to the NCBI Sequence Read Archive (PRJNA1262870). The reading quality was assessed using the NGS-QC Toolkit (version 2.3) (Patel and Jain, 2012). The low-quality reads (Phred score <30; length <70 bases) and adapter sequences were removed to obtain the clean data.

### 2.3 Alignment of reads and gene quantification

The filtered reads were mapped to the *P. cineraria* reference genome (https://zenodo.org/record/7371052#.Y5_1xtJBwfJ) using HISAT2 (version 2.1.0, http://daehwankimlab.github.io/hisat2/) with default parameters. Each sample mapped read was assembled using STRINGTIE (version 2.1.7b, https://ccb.jhu.edu/software/stringtie/) to generate reference-guided assemblies. To identify differential expression genes (DEGs) between heat stress-induced samples and control samples, DESeq 2 (Love et al., 2014), an R package, was used. The p values obtained was adjusted using the p-adjust function (p_adj_) to control the error rate. Genes with a p_adj_ value <0.05 were considered to be differentially expressed. The significance of DEGs was determined by the log2 fold change value being ≥2 (upregulated) or <-2 (downregulated) as the threshold. To cluster DEGs, we performed K-means clustering using R package. Venn diagram was generated with Venny 2.1, an interactive tool for comparing transcripts.

### 2.4 Functional annotation and gene ontology

The identified transcripts in each of the samples were annotated based on homology determined using BLASTx (version 2.2.29+) with threshold e-value <0.005 against publicly available NCBI non-redundant (NR) (https://www.ncbi.nlm.nih.gov/) databases. Annotation of the blast hits against NR was done using the UniProt database (https://www.uniprot.org/). Sequence-based KEGG annotation was done using KAAS (https://www.genome.jp/kegg/kaas/). The DEGs were subjected to BLASTx against the Plant Transcription Factor database (version 5.0 http://planttfdb.gao-lab.org/) to identify transcription factor encoding transcripts with an e-value <0.005. Gene ontology enrichment was performed for the DEGs using the AgriGo v2.0 tool. KEGG pathway enrichment was done using R packages: clusterProfiler (v4.6.2), enrichplot (v1.18.4), and ggplot (v 2_3.4.2) (Wickham, 2016).

### 2.5. Validation of RNA-Seq data by qRT PCR

To validate the data obtained after analyzing the RNA-Seq data, qRT-PCR was performed. The cDNA was synthesized using the BioRad iScript^TM^ cDNA synthesis kit, and the reaction was performed in a BioRad T00 thermal cycler according to the manufacturer’s protocol. Primers for selected genes for qPCR were designed using DNAMAN (Lynnon Biosoft) and Eurofins software following the default parameters. Tubulin β was used as an internal control, with the samples’ biological replicates (n) and technical replicates (N) as 3. The relative expression level of each gene from qRT-PCR data was calculated using the delta-delta method.

## 3. Results

### 3.1 Generation, Mapping, and Assessment of RNA-Seq read

To understand the underlying molecular mechanism of early response to heat stress by *P. cineraria*, cDNA libraries were generated from the leaves of *P. cineraria* grown at 28°C, 45°C (T-45), and 55°C (T-55). The six cDNA libraries were sequenced on the Illumina HiSeq 2X (150) bp platform. In this experiment, approximately 134.5 million paired-end reads were obtained. After removing the adapter sequences and the low-quality reads from the raw data, approximately 120.7 million clean reads were obtained and mapped against the *P. cineraria* reference genome to determine transcriptomic changes during early heat stress. The results related to each group are shown in Table 1. The average GC content of the samples was 45-55%. Principal component analysis (PCA) on RNA-Seq revealed reproducibility and repeatability of the results. The PCA plot indicates a 95% (PC1 = 86% and PC2 = 9%) variation in gene expression datasets with the heat stress conditions (Figure 1), suggesting that heat treatment has significantly disturbed the transcriptome of *P. cineraria* leaf. On average, 87.89% of reads could be mapped to the reference genome, with approximately 80% of them uniquely matched.

**Figure 1:**
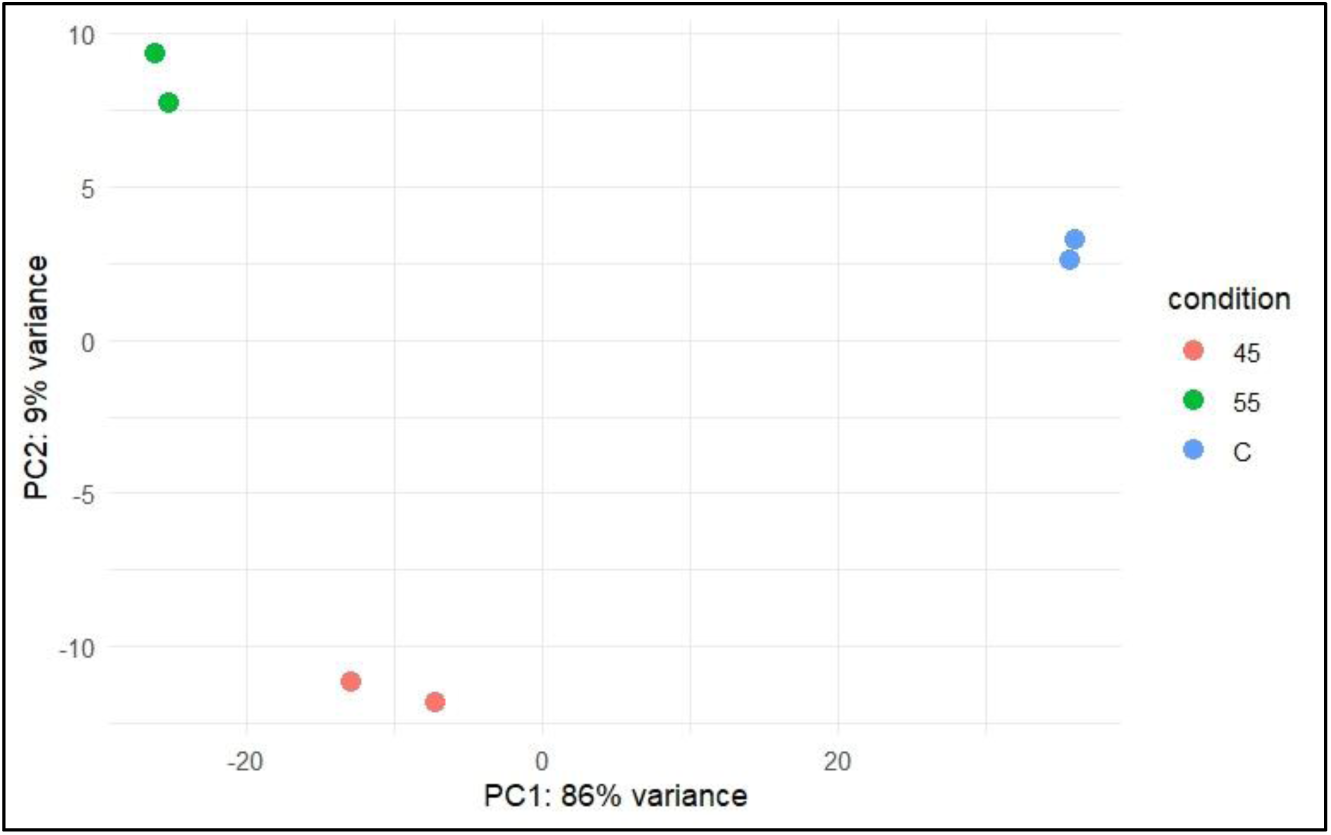
Principal Component Analysis (PCA) of transcriptomic data under different temperature conditions (Control, 45°C, and 55°C). Sample clusters are distinctly based on treatment, with PC1 explaining 86% and PC2 explaining 9% of the total variance. This separation indicates substantial transcriptional reprogramming in response to temperature stress. Each dot represents a biological replicate, and colors denote treatment groups: blue (C) for control, red for 45°C, and green for 55°C.

**Table 1.** Total raw data generated through transcriptome assembly.

| S.No | Sample ID | Treatment | Total Reads | Filtered Reads | Filtered Reads % | Mapped Reads % |
| --- | --- | --- | --- | --- | --- | --- |
| 1 | C-1 | Control | 20124444 | 18261134 | 90.74% | 85.99 |
| 2 | C-2 | Control | 21564048 | 19487707 | 90.37% | 88.72 |
| 3 | 45-1 | Heat | 19850982 | 18021583 | 90.78% | 87.64 |
| 4 | 45-2 | Heat | 22119748 | 19709674 | 89.10% | 87.06 |
| 5 | 55-1 | Heat | 25074012 | 22106510 | 88.17% | 89.58 |
| 6 | 55-2 | Heat | 25806076 | 23205753 | 89.92% | 88.92 |

### 3.2 Differential Expressed Genes in response to heat stress

To identify the DEGs in the T-45 and T-55 samples compared with the control samples, we calculated the transcript abundance of genes with the feature count method and identified the DEGs by setting a threshold of |log2 fold-change| ≥2 & ≤-2 and p_adj_ <0.05. The top 10 significant transcript IDs present in C v/s T-45 and C v/s T-55 are shown in the volcano plot. There were 1151 DEGs (686 up- and 465 down-regulated) in the C v/s T-45, 1562 DEGs (827 up- and 735 down-regulated) in C v/s T-55, and 184 DEGs (123 up- and 61 down-regulated) in T-45 v/s T-55 (Figure 2). The number of DEGs up- or down-regulated in C v/s T-55 samples is more than in C v/s T-45, indicating the T-55 treatment has a pronounced effect. The Venn diagram shows that the DEGs present among C v/s T-45 and C v/s T-55 are 774, C v/s T-45 and T-45 v/s T-55 are 22, and C v/s T-55 and T-45 v/s T-55 are 100; however, 36 DEGs are common between all three samples. The fold change distribution of the DEGs indicates that most and the least number of genes had a fold change of 2-3 and 10-12, respectively (Figure 2E).

**Figure 2.**
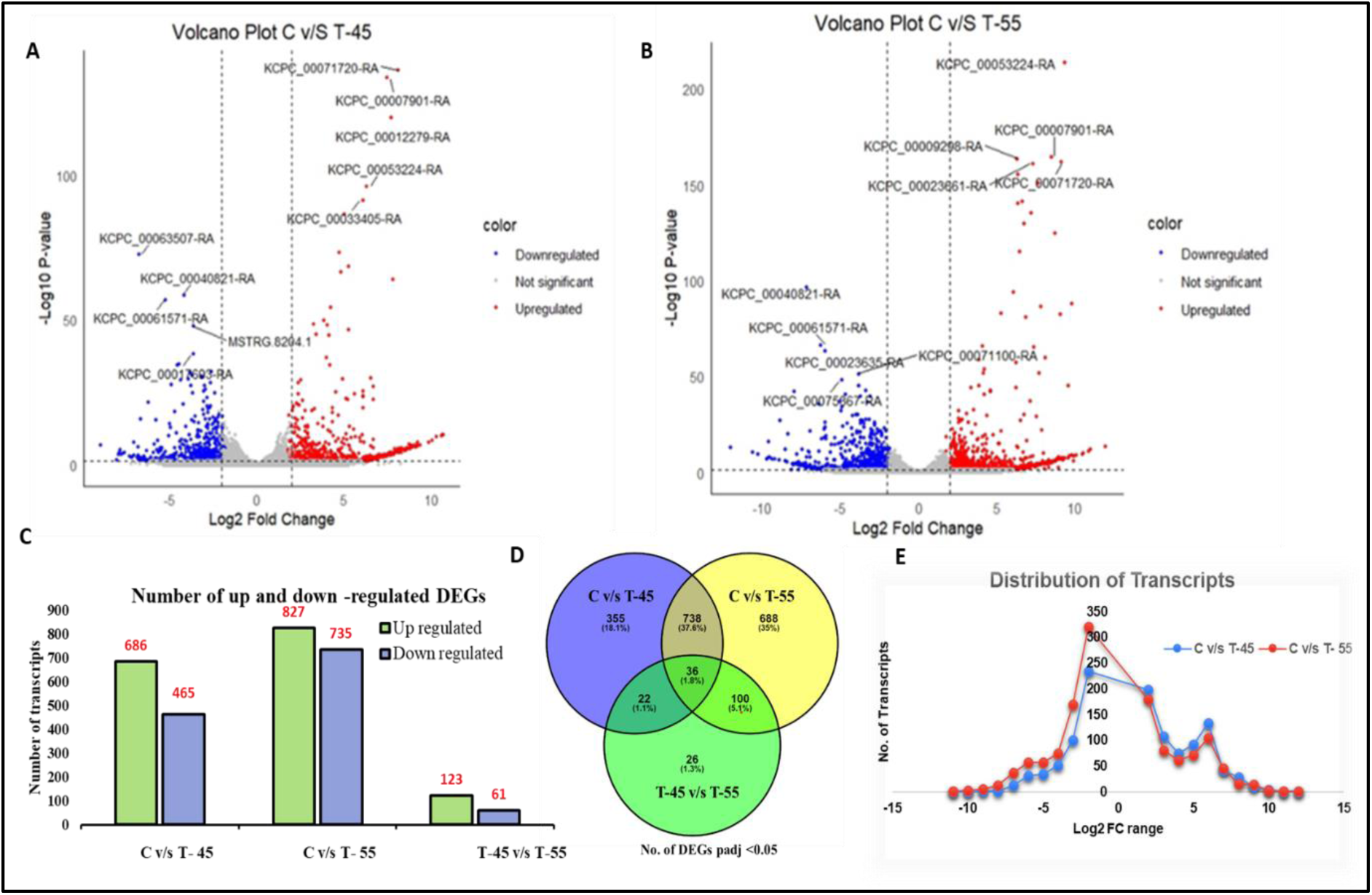
Volcano plots showing differentially expressed genes (DEGs) under heat stress at 45°C and 55°C compared to the control. Each dot represents a gene plotted by its log_2_ fold change (x-axis) and statistical significance as –log_10_ adjusted p-value (y-axis). Red dots indicate significantly upregulated genes, Blue dots indicate significantly downregulated genes, and Gray dots represent genes with no significant change (adjusted p > 0.05 or |log_2_FC| < threshold). The left panel (A) compares control vs. 45°C treatment, while the right panel (B) compares control vs. 55°C treatment. Vertical dashed lines indicate the fold change threshold, and the horizontal dashed line indicates the significance cutoff. Highly significant DEGs are labelled by gene ID. (C) Bar plot shows the number of up- and down-regulated transcripts (y-axis) in each category (C v/s T-45, C v/s T-55, and T-45 v/s T-55) (x-axis). Green bar represents up-regulated and Purple bar represents downregulated genes. (D) Venn diagram shows unique & overlapped DEGs present in each category. (E) The graph represents the distribution of transcripts across the log_2_FC (x-axis) value. Blue dots represent the group of transcripts belonging to the specific range of Log_2_FC in the C v/s T-45 and Red dots in the C v/s T-55 condition.

### 3.3 Expression pattern of DEGs in P. cineraria during heat stress

To study the gene expression patterns in T-45 and T-55 heat stress conditions, the k-means clustering approach was used. All DEGs (1437) between the control and treatment group (T-45 and T-55) following heat stress were grouped in 5 clusters. The distribution of DEGs in each cluster is represented in Figure 3A, and the line plot in Figure 3B represents the expression trend of five clusters (Cluster-1: Red, Cluster-2: Yellow, Cluster-3: Green, Cluster-4: Blue, and Cluster-5: Violet). The expression pattern of individual gene on the basis of Z-score is shown in heat map (Figure 3C). Majority of clustered genes are showing down regulation. Cluster two (128) and three (376) represents the DEGs that were downregulated as stress continues; cluster four (332) and five (626) are highly upregulated at T-45 treatment and their expression dips at T-55; while cluster one (56) DEGs shows exponential increase in expression (Figure 3D).

**Figure 3:**
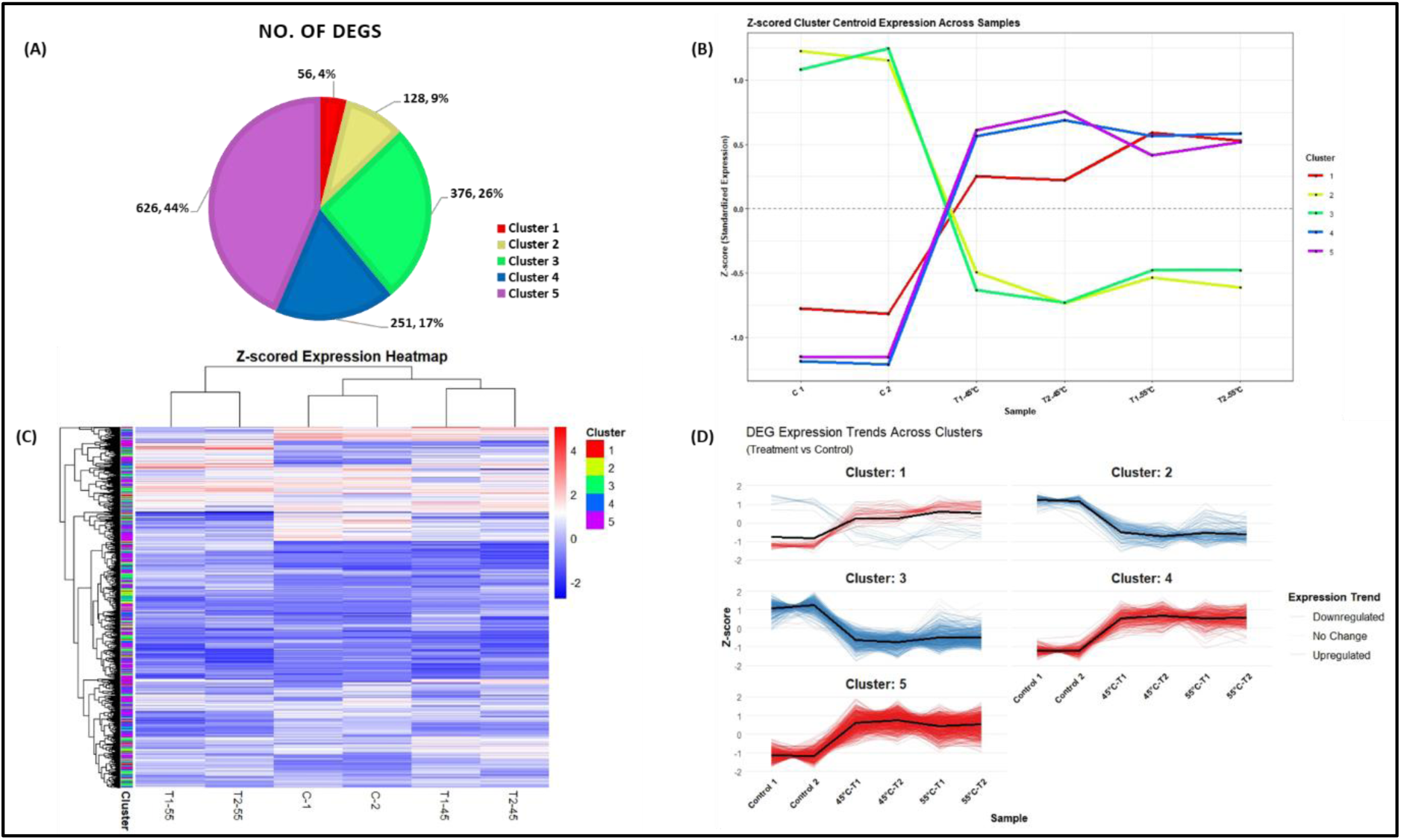
Analysis of DEGs in response to heat stress in *P. cineraria.* (A) k-means clustering of DEGs under heat stress based on gene expression pattern determined by RNA-Seq in five clusters pie chart shows the distribution of DEGs. (B) Z-score cluster centroid expression across samples. The line plot represents the standardised expression profiles of five clusters. Each coloured line corresponds to the centroid of specific cluster. Samples labels for x -axis C: control and T: treatment. The dashed grey line at zero represented the average standardised expression baseline. (C) Heatmap of Z-score of DEGs across five clusters. Rows represent individual genes, and columns represent experimental samples. Gene expression values were scaled to Z-scores to emphasize relative expression pattern. Hierarchical clustering was applied to genes (rows) to reveal expression similarities, while sample clustering is shown on the top. The color bar to the left denotes the K-means cluster. The heatmap color gradient ranges from blue (low expression) to red (high expression). (D) Line plots show the Z-scored expression profiles of DEGs grouped into five clusters based on similar expression patterns across control and heat stress samples. Each line represents the expression trajectory of a gene, and the bold black line denotes the cluster centroid (mean expression trend). Color coding reflects expression trends: red for upregulation, blue for downregulation, and grey lines for genes with subtle changes.

### 3.4 Annotated DEGs Up- and Down-regulated under heat stress

In C v/s T-45 group, annotated DEGs belongs to protein processing in endoplasmic reticulum (ER) pathway, primarily genes that encode for small HSPs including KCPC_00053224-RA (HSP17.9), KCPC_00003282-RA, KCPC_00075853-RA, and KCPC_00013355-RA (HSP17.3), KCPC_00033409-RA, KCPC_00033405-RA, KCPC_00033404-RA (HSP18.5) and KCPC_00041224-RA, KCPC_00071753-RA, KCPC_00013356-RA (SHSP domain-containing protein) that were highly up-regulated. Apart from ER processing, DEGs belong to diverse metabolic pathways. For biosynthesis of secondary metabolites, seven genes were up-regulated and twenty-four were down-regulated. Similarly, two genes for galactose metabolism (KCPC_00006201-RA, and MSTRG.20542.1), and one gene for arginine and proline metabolism (KCPC_00060863-RA) were up-regulated. One gene (KCPC_00046693-RA) and nine other genes in the amino sugar and nucleotide sugar pathway were down-regulated. Importantly, major genes related to plant hormone signal transduction were down-regulated. The details of other DEGs are mentioned in supplementary file 1.

In C v/s T-55 group, annotated DEGs belong to protein processing in ER pathway, that primarily encode for small HSPs, HSP18.5 (KCPC_00069098-RA), HSP17.9 (KCPC_00053224-RA), small HSP 4 (KCPC_00007901-RA), HSP17.3 (KCPC_00009298-RA), in addition to the above-mentioned HSP transcripts. Apart from small HSPs, DEGs belongs to metabolism pathways including biosynthesis of secondary metabolites, where twelve and thirty six genes were up- and down-regulated respectively. Annotated up-regulated genes that were present only in C v/s T-55 condition in response to ROS, are thiamine thiazole synthase (KCPC_00071442), zinc metallopeptidase EGY3 (KCPC_00018764-RA), superoxide dismutase (KCPC_00000985-RA), proline-, glutamic acid and leucine -rich proteins (KCPC_00023440-RA), metallothionein-like protein type2 (KCPC_00068072 and KCPC_00022877) (Figure 4).

**Figure 4:**
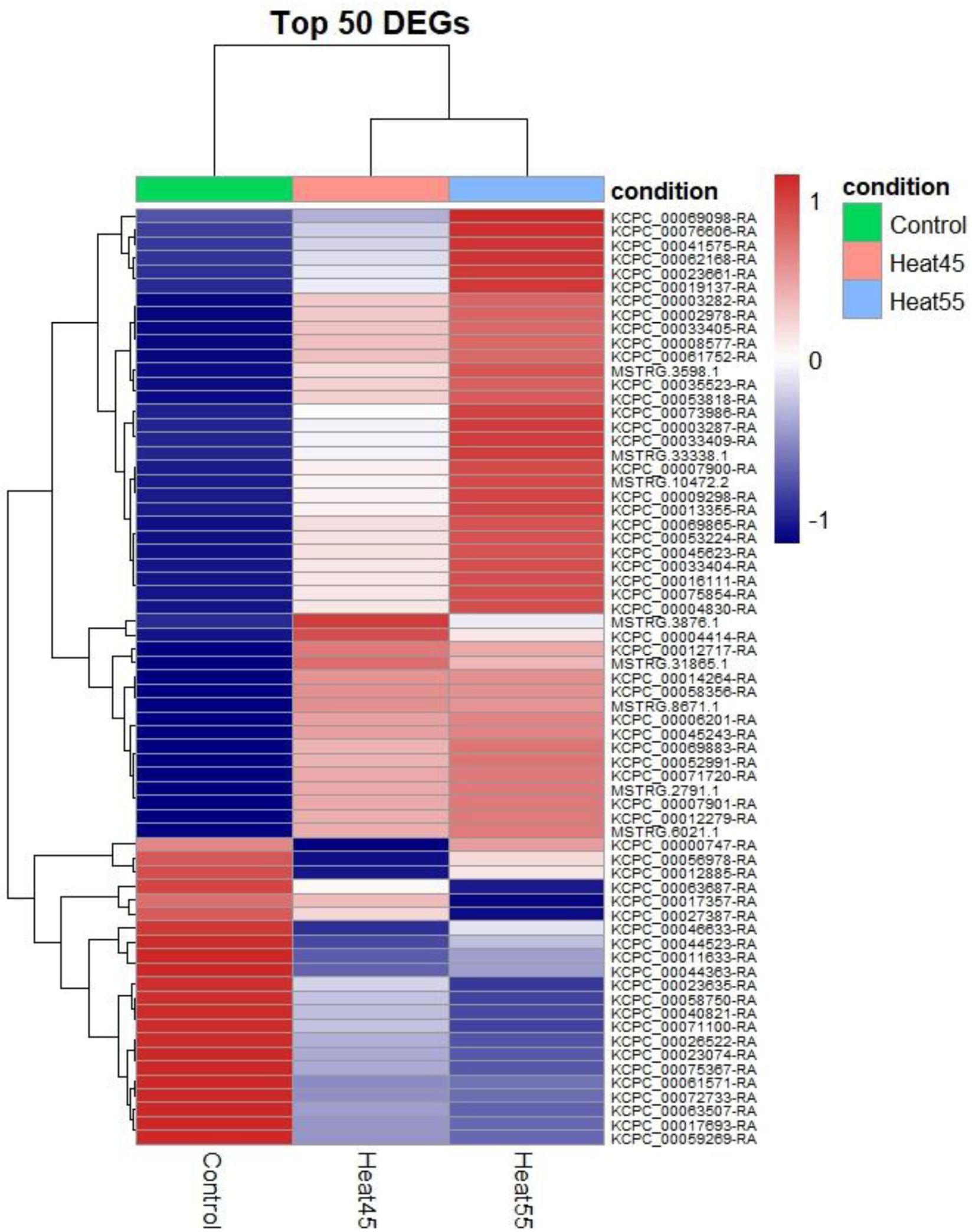
Heatmap showing the expression profiles of the top 50 DEGs, selected based on the lowest adjusted p-values (p_adj_ <0.05) from pairwise comparisons of *Heat45 vs Control* and *Heat55 vs Control* conditions. Rows represent individual genes, and columns represent experimental conditions (Control, Heat45°C, Heat55°C). Color intensity reflects scaled expression (Z-scores), where red indicates high expression and blue indicates low expression relative to each gene’s mean expression across conditions. Clustering was performed on both rows and columns to highlight patterns of co-regulated gene expression under heat stress. Genes: Rows represent individual DEGs (including novel transcripts like MSTRG_ and known genes named as KCPC_).

### 3.5 GO classification of DEGs in P. cineraria

Gene ontology (GO) classification was performed to categorize DEGs into three categories: molecular function (MF), cellular component (CC), and biological process (BP). The GO terms enriched in the C v/s T-45 upregulated DEGs under MF category are unfolded protein binding (GO:0051082), RNA-directed DNA polymerase activity (GO:0003964), and peptidase activity (GO:0070011). Similarly, in the BP category are primary metabolic process (GO:0044238), nitrogen compound metabolic process (GO:0006807), cellular aromatic compound metabolic process (GO:0006725), and protein folding (GO:0006457). In the C v/s T-45 down-regulated genes, GO terms under MF category are predominantly associated with hydrolase activity (GO:0016818), protein heterodimerization activity (GO:0004553), enzyme regulatory activity (GO:0030234), and protein serine/threonine kinase activity (GO:0004674). In BP category, response to biotic stimulus (GO:0009607), carbohydrate metabolic process (GO:0005975), and cell wall organization (GO:0071554) terms were enriched. In CC category, apoplast (GO:0048046), nucleosome (GO:0000786), protein-DNA complex (GO:0032993) were enriched as represented in Figure 5 (A). In C v/s T-55, apart from mentioned in C v/s T-45, up-regulated DEGs were enriched in oxidoreductase activity (GO:0016491) under MF category; response to ethylene (GO:0009723), protein folding (GO:0006457), and phosphorelay signal transduction system (GO:0000160) under BP category; and, mitochondrion (GO:0005739), and cytoplasm (GO:0005737) under CC category. In the C v/s T-55 down-regulated genes, apart from response to biotic stimulus (GO:0009607), and carbohydrate metabolic process (GO:0005975), glucan metabolic process (GO:0006073), and hemicellulose metabolic process (GO:0010410) were enriched in BP category; and, microtubule (GO:0005874) in the CC category. This comparative analysis highlights the dynamic changes in gene function and cellular pathways that occur in response to treatment over time, providing insights into the underlying biological mechanisms affected at each stage.

**Figure 5:**
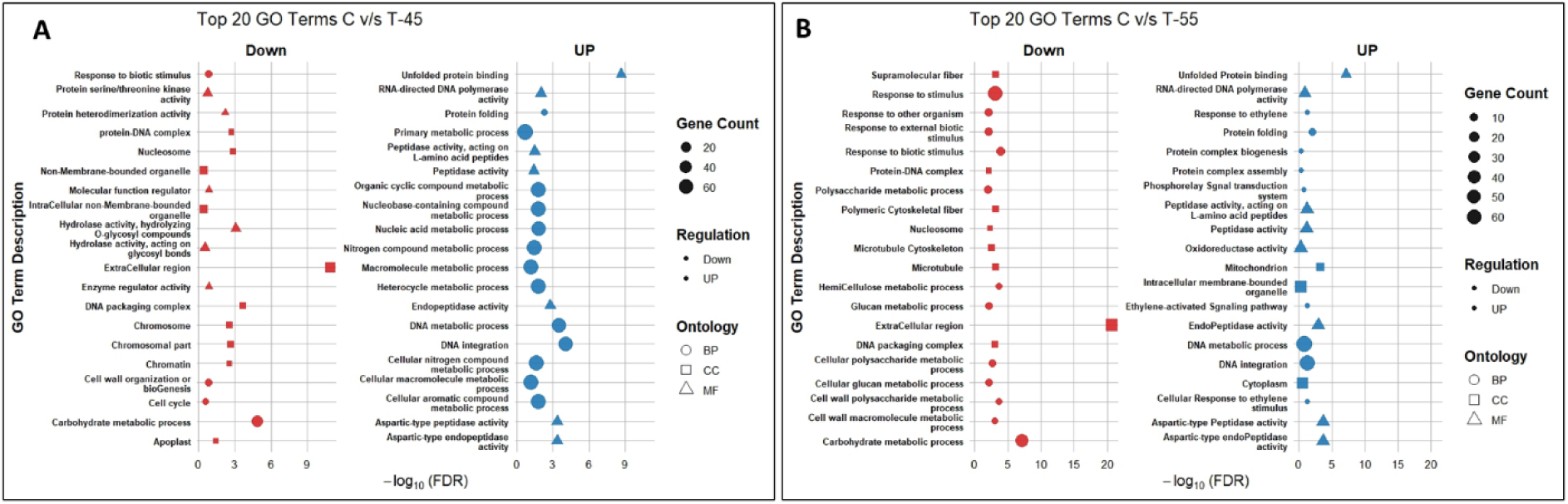
Gene Ontology (GO) enrichment analysis of DEGs under (A) T-45°C and (B) T-55°C heat stress conditions compared to control. Bubble plots display top 20 enriched GO term categories biological process (BP), cellular component (CC), and molecular function (MF) separately for down-regulated and up-regulated genes. The size of each bubble represents the number of genes involved in each GO term, and the use of different shapes distinguishes between BP, CC, and MF categories. The x-axis represents the statistical significance (–log_10_ FDR), and the y-axis lists enriched GO terms.

### 3.6 KEGG annotations of DEGs under extreme heat stress

In different heat conditions, enriched KEGG pathways are shown in the figure 6. In C v/s T-45 condition, protein processing in the ER (ko04141) was assigned to 27 DEGs. This is followed by amino sugar and nucleotide sugar metabolism (ko00520), linoleic acid metabolism (ko00591), alpha-linolenic acid metabolism (ko00592), and zeatin biosynthesis (ko00908), which were assigned to 12, 6, 6, and 4 DEGs, respectively. However, in C v/s T-55, protein processing in ER was assigned to 35 DEGs, followed by phenylpropanoid biosynthesis (ko00940), amino sugar and nucleotide sugar metabolism (ko00520), and flavonoid biosynthesis (ko00941), which were assigned to 14, 14, and 7 DEGs, respectively. In the T-45 v/s T-55 condition, protein processing in the ER (ko04141) was assigned to 18 DEGs, followed by the MAPK signaling pathway (ko04016), endocytosis (ko04144), and Linoleic acid metabolism (ko00591), which were assigned to 5, 5, 2 DEGs respectively (Figure 6). In KEGG pathway enrichment analysis, protein processing in the ER (ko04141) was most significantly enriched, indicating that it plays a central role in *P. cineraria* under heat stress.

**Figure 6:**
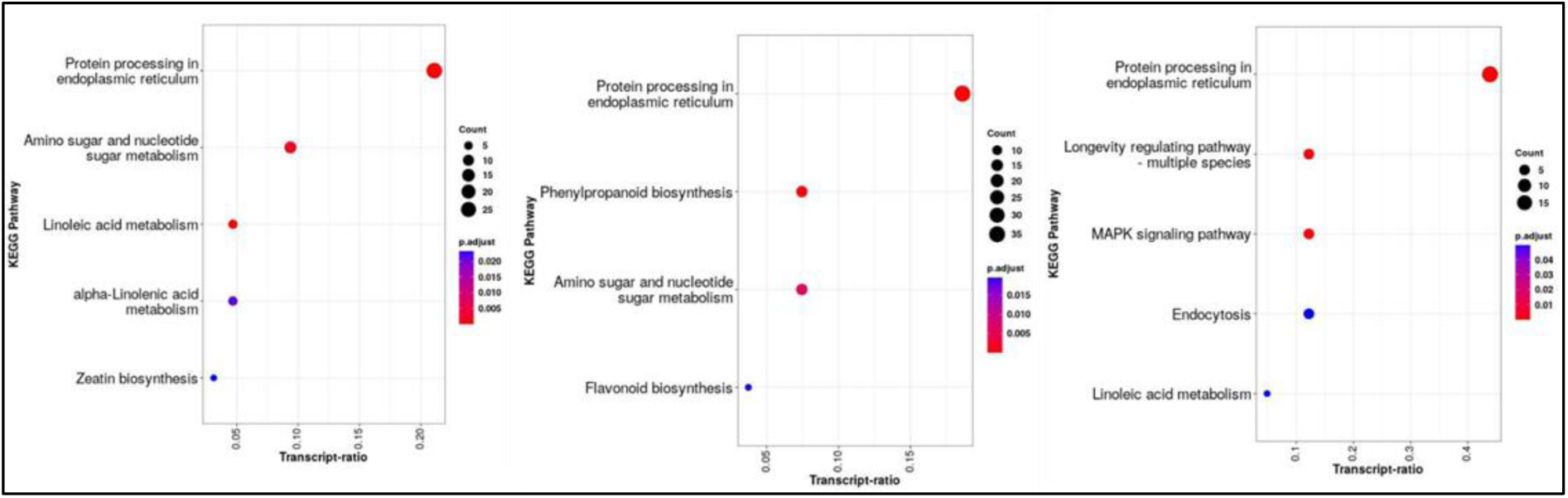
KEGG pathway enrichment analysis of DEGs under varying heat stress conditions. The three panels represent the pathway enrichment for DEGs under different treatments: Left: C v/s. T-45; Middle: C v/s. T-55; Right: T-45 v/s T-55. Each bubble represents a KEGG pathway: x -axis: Transcript ratio (proportion of DEGs mapped to the pathway), y-axis: KEGG pathway name. Color gradient: Adjusted p-value (p_adj_), indicating statistical significance (more red = more significant). Dot size: Number of DEGs (Count) associated with each pathway.

### 3.7 Differential expression of transcription factor (TF) families

In plants, TFs, such as WRKY, ERF, MYB, HSF, NACs, ARFs families are known to be differentially expressed under abiotic stress (Liu et al., 2024). In *P. cineraria*, a total of 866 transcripts were predicted to belong to 50 different TFs families. Among these, 134 transcripts belong to nuclear factor Y (NF-Y) family, which is a heterotrimeric complex of NF-YA, NF-YB and NF-YC. During heat stress, the NF-YA (95), NF-YB (33), and NF-YC (6) transcripts were highly differentially expressed and majority of them were up-regulated in both C v/s T-45 and C v/s T-55 conditions. Other important transcription families such as MYB (86), bHLH (71), NAC (70), ERF (68), WRKY (52), HSF (21), and bZIP (19) were differentially regulated under heat stress. On the basis of the log_2_FC value, the top 20 transcripts of each family are represented in Figure 7.

**Figure 7:**
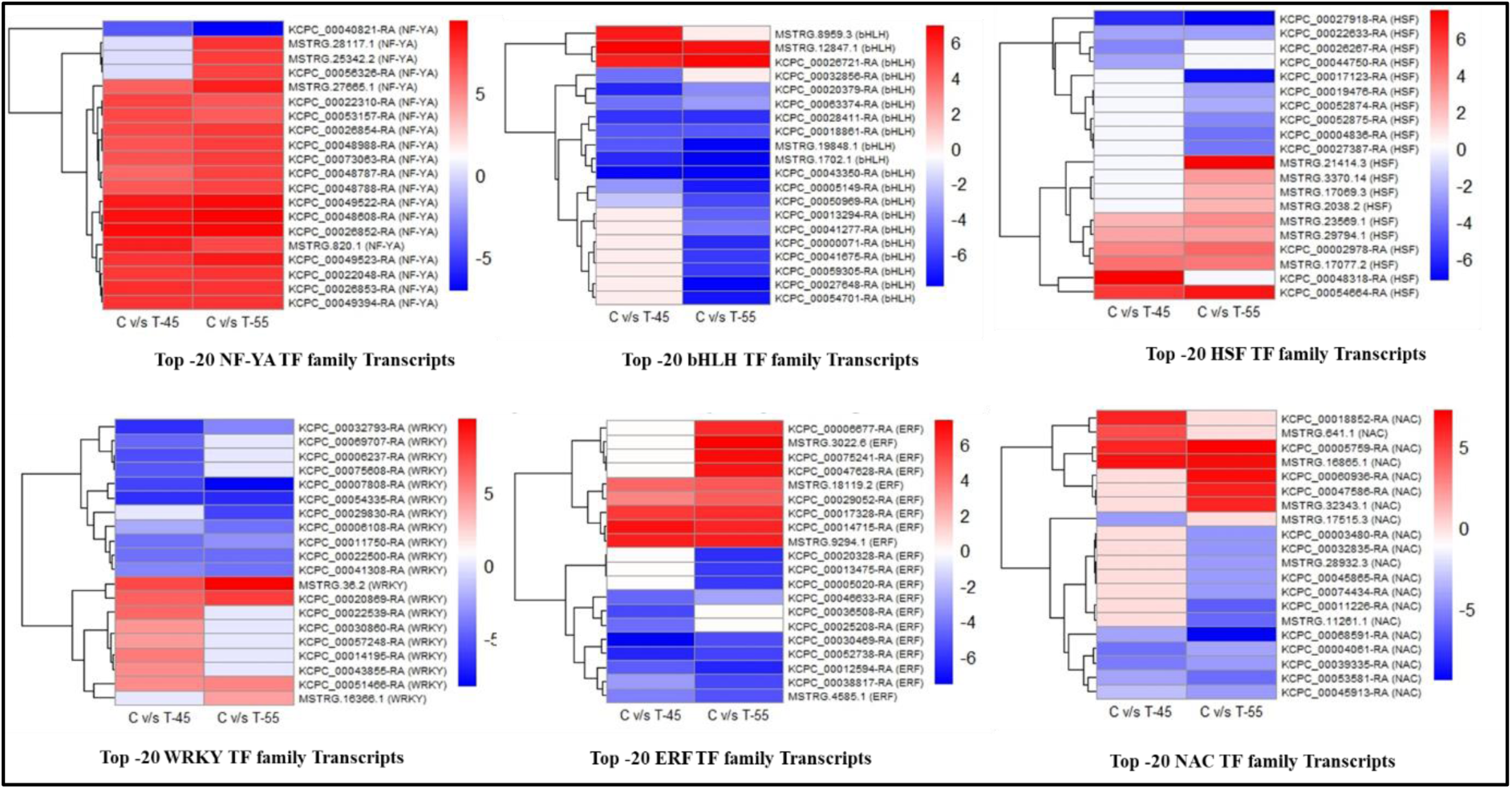
Heatmap visualization of top differentially expressed transcripts across six transcription factor (TF) families (NF-YA, BHLH, HSF, WRKY, ERF, and NAC) under control (C) and stress (T-45) conditions. Each heatmap shows the top 20 transcripts selected based on highest log_2_ fold change (log_2_FC). Color gradients represent log_2_FC values: red indicates up-regulation, blue indicates down-regulation, and white represents no change. Rows represent individual transcripts; columns represent experimental conditions. Clustering is based on Euclidean distance and complete linkage for rows.

### 3.8 Validation of DEGs by quantitative real-time PCR

The validation of a few differentially expressed transcripts was done by quantitative real-time PCR. The representative genes including a few heat shock proteins and heat shock transcription factors with significant changes were examined using gene-specific primers. All the transcripts showed a similar expression pattern observed in bioinformatic differential analysis results from high throughput sequencing (Figure 8).

**Figure 8:**
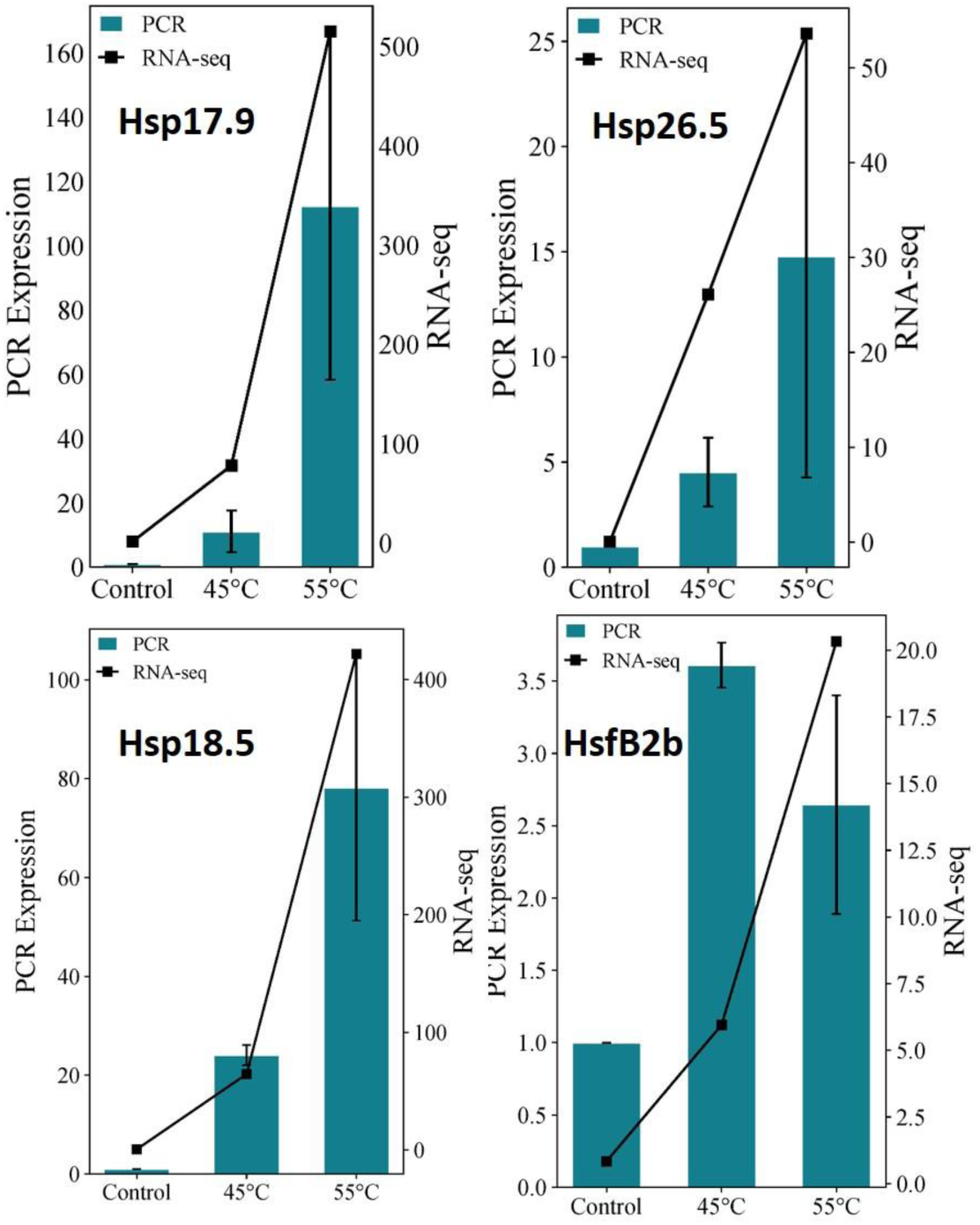
Bar plots represent quantitative PCR (qPCR) expression levels (left y-axis, teal bars) and RNA-Seq transcript abundance in FPKM (Fragments Per Kilobase of transcript per Million) mapped reads; right y-axis, black lines with square markers) for selected heat-responsive genes—Hsp17.9, Hsp18.5, Hsp26.5, and HsfB2b subjected to control (28°C), 45 °C, and 55 °C heat stress conditions. qPCR data are shown as mean ± standard error from three biological replicates, and using reference gene Tubulin β.

## 4. Discussion

Temperature is an important environmental factor, involved in regulating plant growth and development, and rise in temperature beyond threshold impacts the crop productivity. *P. cineraria* is a versatile legume tree, inhabiting abundantly in arid and semi-arid regions of India. It can withstand wide range of temperature > 45°C (Sasi et al., 2024). However, the genes involved in regulating heat stress have not been studied extensively. This study identified 1151 and 1562 DEGs under 45 °C and 55 °C heat stress, respectively, in *P. cineraria* using RNA-Seq analysis. There are many promising candidate genes from multiple families that are potentially involved in heat stress tolerance of *P. cineraria* are discussed here.

### 4.1 Transcription factors

When plants experience heat stress, they immediately initiate the expression of heat shock transcription factors (HSFs), especially the HSFA family, which acts as master regulators and enhances the expression of downstream genes like *HSPs*. The role of HSFs during stress conditions is well understood and explored (Bakery et al., 2024; Guo et al., 2016). *AhHSF3, AhHSF28* in peanut and *AtHSFA1d* in Arabidopsis induce during drought conditions and enhance drought resistance (Wang et al., 2023). However, HSFB1 and HSFB2b are known to negatively regulate the expression of some *HSFs* and *HSPs* (Ma et al., 2022). But knockout of *hsfb1* and *hsfb2b* leads to lower thermotolerance than wild type, indicating that they might promote the activity of HSFA1 by repressing specific *HSPs* under heat stress conditions (Guo et al., 2016). Here, the *HSFB-2a* (KCPC_00002978) gene belonging to cluster 4, is highly upregulated at T-55, suggesting it might be involved in heat response. Apart from HSFs, APETALA2 / ethylene responsive factor (AP2/ERF) is one of the largest transcription factor families featured for the conserved AP2/ERF DNA binding domain and regulates the expression of the heat-inducible genes (Ma et al., 2024). ERF and DREB family genes are expressed in wheat during heat stress in both tolerant and susceptible cultivars. However, in tolerant cultivars, some AP2 genes were up-regulated (Magar et al., 2022). In *P. cineraria*, seven genes of the AP2/ERF family were highly upregulated, and eight were down-regulated, indicating the role of specific APS/ERF family in regulating heat response. Squamosa promoter-binding-like protein (SPL) is a class of transcription factors (TFs) unique to plants and regulates the growth phase transitions. SPL genes have been extensively characterized from various species. The expression level of specific SPL genes is up- and down-regulated during stress conditions and regulates the activities of SOD and peroxidase (Hou et al., 2018). When *Medicago sativa* is subjected to salt stress, the expression of specific *MsSPL* sharply increases (Ma et al., 2022). At T-55 in *P. cineraria*, the transcript (MSTRG.26639.1) encoding for SPL shows Log_2_Fold change of 7.03 and the expression of peroxidase (KCPC_00064340-RA & KCPC_00016470-RA) and SOD transcript (KCPC_00000985) was upregulated.

The photoperiod pathways are regulated by CONSTANS (CO), which upregulates the expression of downstream genes *FLOWERING TIME (FT)* and *SUPPRESSOR OF OVEREXPRESSION OF CO (SOC1)*. *CO* is known as B-box protein, a zinc finger binding domain. The role of B-box in regulating flowering time is well studied. However, the role of B-box protein during abiotic stress has not been explored fully. In Arabidopsis and banana, the transcripts of distinct BBX increase in response to chilling. In Arabidopsis, the overexpression of BBX-24 confers salt tolerance (Nagaoka & Takano, 2003). Yang et al., (2014) elucidated the involvement of BBX-24 in drought and low temperature in *Chrysanthemum morifolium* through gibberellin acid pathways. During heat stress at T-45 in *P. cineraria*, the transcript level of zinc finger protein CONSTANS -Like 2 (KCPC_00015115) is highly upregulated, and at T-55, another zinc finger protein CONSTANS -Like 13 (KCPC_00027732) is found to be up-regulated. Other transcription factors like basic bHLH, NAC, WRKY, and basic leucine zipper were downregulated.

### 4.2 Transporters

Nitrate transporter 1 (NRT) / peptide transporter (PTR) family (NPF) proteins are well known for transporting nitrate or di/tripeptides. Recently, the NPF family of proteins has been found to participate in transporting plant hormones. Among the NPF protein family, NRT1.1 has multiple functions and is involved in abiotic stress resistance (Fang et al., 2021). In *P. cineraria*, the expression of *NRT4.6* (KCPC_00059882) and 5.6 LIKE (KCPC_0004141) transcript was up-regulated, suggesting their role in transporting abscisic acid. However, the role of NRT 5.6 like is not been fully explored yet. Moreover, two transcripts related to the NPF family (KCPC_00057278 and KCPC_00057277) were downregulated.

Polyol transporters (PLTs) belong to the monosaccharide transporters family, involving transportation of polyols and monosaccharides. In *Lotus japonicus,* the *LjPLT1/ 3* and *9* were differentially expressed during salinity and osmotic stress conditions (Tian et al., 2017). Similarly, *OeMaT1* was significantly induced in response to salinity and drought stress in *Olea europaea* (Conde et al., 2011), and *OsPLT3* was up-regulated during heat stress in rice (Kong et al., 2020). In our data, the expression of PLT3 (KCPC_00044566) is up-regulated at both conditions.

Organic cation/carnitine transporter (OCTs) is a less explored plant transporter class. They are characterized as carnitine transporter in the plasma membrane. Küfner and Koch (2008) studied the expression of the Arabidopsis OCT family during drought, salinity, and cold stress. It was found that during all three stresses, the *AtOCT5* responds strongly, whereas *AtOCT3* expression is induced only during cold treatment (Küfner & Koch, 2008). However, no reports have yet shown the participation of OCT3 during heat stress. *In P. cineraria,* (KCPC_00051806) *OCT3 LIKE* is up-regulated at T-45, however, during T-55, the expression of *OCT3* (KCPC_ 00068993) further increases to log_2_ fold change of 6.

Mitochondrial carrier proteins (MCs) are important in transporting molecules to and fro from cytosol. In Arabidopsis, basic amino acid carriers 1 and 2 (BAC1 and BAC2) transport basic amino acids like lysine, arginine, histidine, and ornithine (Monné et al., 2019). Hyperosmotic stress mainly regulates the expression of the *BAC2* gene, which leads to proline accumulation (Toka et al., 2010). Interestingly, during heat stress, the *BAC2* (KCPC_00026262) was highly up-regulated in both conditions in *P. cineraria*.

### 4.3 Cell wall remodeling and carbohydrate metabolism

Cell wall remodeling during stress is an important event. Various enzymes like endo-1,4-β-D-glucanase (EGase), xyloglucan endo-β-transglucosylases/hydrolases (XTH), pectin methylesterase (PME), polygalacturonase (PG), pectin/pectate lyase-like (PLL), Fasciclin-like arabinogalactans (FLAs) and pectin acetylesterase (PA) play a key role in the modulating cell wall. In our data, 44 transcripts are involved in cell wall modification; among those transcripts, 42 are highly down-regulated, and two transcripts are up-regulated (KCPC_00065026-RA and KCPC_00051276-RA). The family of genes involved in the strengthening of cell wall are XTH and FLAs. Numerous studies have shown the roles of the XTH family in various environmental stresses. In pepper, *CaXTH3* can be induced by cold, salt, and drought stress (Choi et al., 2011). Similarly, the expression of *MtXTH13* was upregulated during salt stress in *Medicago truncatula* (Xuan et al. 2016). However, some isoform genes of XTHs are found to be down-regulated. During drought, the expression of *AtXTH* 6, 9, 15 and 16 was downregulated in Arabidopsis (Clauw et al., 2015) and the expression of *SeXTH 4/5,7/8, 9, 31,* and *32* was down-regulated in *Salicornia europae* L. (Tiika et al., 2021). In pearl millet root, XTH was down-regulated during heat stress (Singh et al., 2024). In our data, transcripts encoding for XTHs belonging to cluster 3 (KCPC_00045913, KCPC_00015705, and KCPC_00059319) are significantly downregulated during the C v/s T-55 condition.

Fasciclin-like arabinogalactans (FLAs) are involved in biosynthesis and strengthening the cell wall. During drought in Arabidopsis, *AtFLA2* and *AtFLA9* were downregulated (Clauw et al., 2015). The transcripts encoding for FLA1 (KCPC_00009175), FLA11 (KCPC_00004836 and KCPC_00019476) and FLA12 (KCPC_00027387 and, KCPC_00052875) were also downregulated and FLA3 (KCPC_00065026) was upregulated. The expansin (*EXP*) gene family is involved in cell wall loosening, and during heat stress, many *EXPs* were upregulated in potato, whereas *EXPB2* was highly downregulated (Chen et al., 2019). We observed at the T-55 condition that EXP (KCPC_00008414 and KCPC_00000071) was highly downregulated.

The chitinases are differentially regulated during drought, salt, low and high temperatures (Zhou et al., 2020). During temperature fluctuations, chitinase gene expression is reported to be down-regulated in maize, wheat and grape (Lamba et al., 2022; Liu et al., 2012; Wang et al., 2024). Here, 19 chitinase genes are down-regulated, and only two genes (KCPC_00009238 and KCPC_00046693-RA) are upregulated.

#### β-mannanase

Abiotic stress affects the carbohydrate metabolism. The transcripts related to the specific carbohydrate metabolism pathway, i.e., mannan endo-1,4 - beta-mannosidase (β-mannanase), were highly up-regulated during salt stress in two contrasting genotypes of *Carex rigescens* (Zhang et al., 2020). β-mannanase is involved in the degradation of mannan polymers via random cleavage of the β-d-1,4-mannopyranosyl linkage. Mannan polymers are one of the important components of hemicellulose fractions in plants (Rahmani et al., 2017). In another study, transcriptomic data of a 2-day-old maize seedling showed that during heat stress of 42°C, genes involved in the biosynthesis of carbohydrates were upregulated (Li & Ye, 2022). A similar trend is observed in our data, and *β-mannanase* gene was significantly upregulated in both conditions. Their exact role during stress conditions has not yet been explored.

### 4.4 Cellular signaling

#### Reactive oxygen species (ROS)

ROS are toxic by-products of cellular metabolism and the level of ROS is elicited during different stresses, including heat stress, and causes oxidative damage. Organelles like peroxisomes, where ROS is generated and detoxified, are involved in metabolic pathways against oxidative stress. We found one gene related to peroxisome, Acetate/butyrate-CoA ligase AAE7 (peroxisomal) (MSTRG.2450.1) with log_2_FC of 6.5 belongs to cluster 5. The transcript level of AAE7 during T-45 and T-55 seems to be consistent and high. The beta-oxidation of very long fatty acids (VLFAs) occurs in peroxisomes, which cannot be oxidized by mitochondria. VFLAs are converted into acylcarnitines and translocate to mitochondria via carnitine translocase. In mitochondria, acylcarnitines are converted to acyl-coenzyme A, which enter into the citric acid cycle. In our data at both temperatures, the expression of KCPC_00051806-RA transcript which encode for organic cation/carnitine transporter is also high. However, its log_2_FC value is lesser at T-55 than T-45. The following results suggest that during heat stress, lipid mobilization increases and help a plant to prevent carbon loss during lipid mobilization, as plant is undergoing stress conditions.

During stress conditions, chloroplast triplet chlorophylls and electron transport chains are the sites of ROS production. In chloroplasts, pseudoproteases like Ethylene-dependent Gravitropism-deficient and Yellow-green (EGY3) are present and perform the proteolytic cleavage of proteins anchored to the cell membrane. The expression of *EGY3* during heat and salt stress increases, as shown in *Vigna unguiculata* (Selinga et al., 2022). Similar results were obtained in Arabidopsis when exposed to high temperatures (Adamiec et al., 2022). In our data, the expression of *EGY3* (KCPC_00018764) belongs to cluster 5 was significantly up-regulated compared to the control in T-55. The transcript abundance of the *EGY3* gene increases not only during heat stress but also in response to salt stress, as observed in Arabidopsis (Zhuang et al., 2021).

#### Ferredoxin

The photosynthesis process is highly sensitive to high temperatures as heat stress directly affects the photosynthetic apparatus present in chloroplast. Damage to the photosynthetic apparatus leads to the generation of ROS. Ferredoxins (FDXs) are soluble iron-sulfur proteins involved in the ROS scavenging system to reduce the toxicity of ROS by regenerating ascorbate. During heat stress, it is observed that in *P. cineraria*, the transcript level of FDXs is higher at T-45 (MSTRG.26370.1) than at the T-55 condition. This suggests that high transcript levels of FDXs prevent the effects of excessive ROS production by protecting cells from damage and helping plants strive at high temperatures. However, a decline in the expression of FDXs at T-55C, is possibly due to the destabilization of FDX transcripts. Using overexpression studies of specific types of FDXs helps plants like tomatoes to combat heat stress (Meng et al., 2022), as well as in rice, against salinity stress (Huang et al., 2020). Similar results were obtained via knockdown studies in Arabidopsis; at 44℃ heat stress, there was an increase in the reduced form of ascorbate via FDXs (Lin et al., 2015). Interestingly, in tall fescue, the abundance of an FDXs gene was increased during heat stress in heat tolerant PI 578718 cultivar (Hu et al., 2014). Another study showed the high abundance of FDX transcripts in ryegrass during basal and acquired thermotolerance (Chen et al., 2023).

#### Glutathione S-transferases (GSTs)

are a large, ubiquitous, and complex family of enzymes that regulate plant growth, development, and stress response. During metabolic processes, distinct forms of toxic components are produced. The level of toxic compounds is alleviated during stress conditions (Gao et al., 2020). In such a situation, initially, cytochrome P450s convert the toxic compounds into reactive electrophiles. GSTs reduce the toxicity of reactive compounds and increase their solubility, which will be targeted for final degradation (Wang et al., 2019). The GSTs are classified into fourteen classes in plants. In plants, phi and tau GSTs correlate highly with stress tolerance (Islam et al., 2019). As observed in overexpression studies of Grape *GST13* in Arabidopsis, it increases tolerance towards salinity, drought, and methyl viologen stresses (Xu et al., 2018). Another report shows that the overexpression of *TaGSTU1B* and *TaGSTF6* genes in wheat enhances tolerance towards drought stress (Gallé et al., 2009). In our data, six DEGs are annotated as GST. At T-55, GST (KCPC_00031464, KCPC_00022778, and KCPC_00022780) log_2_FC is 2 and KCPC_0004706 is downregulated. This indicates a specific GST gene’s role in buffering redox and managing stress. However, at T-45, GST genes are downregulated only.

#### Cytochrome

Plant Cytochrome P450s (CYPs) monooxygenases are one of the largest oxidoreductase protein families involved in crucial biosynthetic reactions. It is well known that specific *CYP* genes are also involved in plant stress response (Pandian et al., 2020). *CYP* genes that participate in flavonoid production during heat stress are differentially expressed. In *Lolium perenne* and *Festuca arundinacea*, *CYP73A, CYP75A*, and *CYP75B* genes were highly upregulated under temperature stress (Tao et al., 2017). Whereas, in the case of *Rhazya stricta* during heat stress, transcripts encoding for CYP were highly downregulated (Obaid et al., 2016). In this study, the abundance of *CYP 450 787A-like*, with respect to control, is found to be significantly high in both T-45 and T-55 under heat stress. A total of eighteen genes were annotated for CYPs. At T-45, three unique *CYP* genes (KCPC_00065860, KCPC_00059992, and KCPC_00060330) were up-regulated, and one KCPC_00071935 gene was downregulated. Meanwhile, at T-55, eleven genes were upregulated and four downregulated. The *CYP78* family generally stimulates plant growth and regulates plant architecture. The reports related to the *CYP78* family during abiotic stress are limited. Recently, the expression of *CYP78A7a* in eggplant was upregulated during salt stress and downregulated during dehydration stress (Shen et al., 2024). Another important class of *CYP* expressed during stress is *CYP707A*. It encodes for Abscisic acid 8’-hydroxylase (ABA 8’-hydroxylase). This enzyme is involved in the catabolism of plant hormone ABA via the oxidative pathway. In Arabidopsis, high temperatures trigger the subtle expression of specific *CYP707A* (*CYP707A1*, *CYP707A4*) genes (Baron et al., 2012). The transcript of *Abscisic acid 8’-hydroxylase 1* was predominantly down-regulated in cucumber under heat stress (Wang et al., 2020). However, in our data, the relative abundance of *Abscisic acid 8’-hydroxylase 4-like* isoform X2 during heat stress is quite high compared to control. The possible reason for the high abundance is to maintain homeostasis of abscisic acid; however, it needs further validation.

### 4.5 Molecular chaperones

The accumulation of misfolded, unfolded, or denatured proteins in the ER under abiotic stress leads to ER stress. After high-temperature stress in maize and garlic (Yang et al., 2024), the ER protein processing was significantly upregulated, consistent with our results. At high temperatures in *P. cineraria,* KEGG enrichment analysis showed that protein processing in the ER (ko04141) pathway is most enriched including 35 transcripts, suggesting that the ER is experiencing stress and triggers the unfolded proteins response (UPR), which might play an essential central role during stress.

HSPs are known to be expressed during the initial stage of stress. In this study, 32 DEGs were annotated as HSPs. The majority of expressed HSPs belong to the small HSPs family (HSP17.3, HSP17.7, HSP18.5, HSP17.9, and HSP22); however, 7 HSPs belong to higher HSPs i.e. HSP83 (MSTRG.13111.2), HSP70 (KCPC_00041575, MSTRG.18593.2 and MSTRG.25456.1), HSP60 (MSTRG.17105.1) and HSP 40 (KCPC_00022608 and KCPC_00060936). This finding is similar to *Clematis apiifolia* and *Nelumbo adans* under heat stress (Gao et al., 2017; Liu et al., 2016). Apart from molecular chaperones, activators of HSP90 ATPase activity (AHA) (KCPC_00016111 and KCPC_00005233) are also up-regulated. Recent studies have shown the interaction between HSP70 and Bcl-2-athanogene (BAG) proteins under heat stress. In Arabidopsis, *AtBAG6* is significantly up-regulated during heat stress (Doukhanina et al., 2006). Similarly, one gene annotated for BAG (KCPC_00034789) is identified as a heat-inducible gene. However, its interaction with specific HSP70 requires further experiments.

The DEAD-box RNA helicase (RH) is known to act as RNA chaperones. The expression of DEAD-box RHs is altered by high temperature, salinity, and light intensity. During heat stress in tomato, the expression of *SlDEAD31* was up-regulated, and *SlDEAD30* was downregulated. Moreover, tomato plants overexpressing *SlDEAD31* could tolerate salt and drought stress (Zhu et al., 2015). Similar studies were carried out in rice, *OsBIRH1* overexpressing rice lines showed enhanced tolerance towards oxidative stress and disease resistance (Li et al., 2008). Recently, the expression of *Brassica rapa* (*BrRH22*), a chloroplast targeted DEAD-box RH during heat stress, was upregulated 4-fold. Arabidopsis over-expressing *BrRH22* plants grew better than control under high drought and salinity stress (Nawaz et al., 2018). In our data at T-55, the expression of DEAD-box ATP-dependent RH 38 (KCPC_00016348-RA) is highly up-regulated, and DEAD-box RHs are known to regulate the expression of dehydration positively -responsive element binding (DREB) transcription factor. Interestingly, the expression of DREB protein 2C (KCPC_00064317-RA) in T-55 is also up-regulated, possibly due to the helicase and contributing to tolerance towards stress.

### 4.6 Ubiquitin-Proteasome System

In all eukaryotes, the ubiquitin-proteasome system (UPS) plays an important role in the selective degradation of intercellular proteins. The ubiquitin system has three enzymes: the E1-activating enzymes, the E2-conjugating enzymes, and the E3-ligases, that lead to the degradation of target proteins. The F-box subunit of the Skp1-Cullin -F-box (SCF) E3 ligase is involved in abiotic stress response by interacting with multiple substrate domains. Hence, F-box proteins (FBPs) are one of the plants’ most diversified and largest superfamilies. They are known to play essential roles in regulating protein homeostasis during plant development and stress. The change in expression of mRNA transcripts of FBS during abiotic stress is reported in many studies. Like in sweet potato, the expression of FBX75, FBX208, and FBX232 transcripts were significantly high during salt stress, and the FBX130, FBX 65, FBX 127, and FBX220 transcripts were high during drought stress (Amoanimaa-Dede et al., 2022). The pepper F-box protein transcript was up-regulated after cold and salt treatment (Chen et al., 2014). Likewise, *Phaseolus vulgaris* F-box transcripts accumulate in response to osmotic stress (Maldonado-Calderón et al., 2012). The overexpression of the wheat TaFBA1 in tobacco showed increased tolerance against heat stress (Li et al., 2018). However, overexpression of *Glycine max GmFBL144* significantly reduced the Arabidopsis drought tolerance (Xu et al., 2022). Similar results were obtained in rice overexpressing the F-box gene, which was more sensitive to drought conditions than wild type (Yan et al., 2011). However, the presence of the lysin motif (LysM) and F-box domain in a single protein suggests its role in the degradation of LysM-interacting proteins. The role of LysM/F-box - containing proteins in plants during stress has not been studied yet. Recently, the function of the Arabidopsis LysM/F-box containing proteins has been elucidated in normal conditions. These proteins fine-tune glycine metabolism (Guo et al., 2021). In our transcriptomic data at T-45, the transcript of F-box protein At1g55000 isoform X1 (KCPC_00067447) is significantly upregulated by a 7.4-fold change. Elevation in the transcript under heat stress has not yet been explored.

## 5. Conclusion

*Prosopis cineraria* is a highly thermotolerant plant, and grows well in all types of climatic constraints which is evidenced by the fact that new foliar growth, flowering and fruiting occur during dry and hot months when most other trees of the desert remain leafless or dormant. Thus, *P. cineraria* represent a repository of genes for heat stress tolerance, which are likely to be constitutively expressed and can be utilized for the genetic improvement of crops. RNA-Seq based genome-wide analysis gives complete transcriptome profile of differentially expressed genes and this is most recent technique available to study gene family annotations. Our focus is on heat stress responsive genes in *P. cineraria* while it is now evident from this study that these genes also play role in multiple abiotic stresses. The involvement of multiple gene families and their complex regulation for heat stress tolerance in *P. cineraria* will enable us to understand the mechanism of stress tolerance in arid plants.

## Supporting information

Supplementary file 1 : P. cineraria DEGs

## Acknowledgements

HS thank BITS Pilani for the Institute fellowship. MJ acknowledges the Department of Biotechnology, Government of India, for financial support with Ramalingaswami Re-Entry Fellowship award (Ref. No. BT/RLF/Re-entry/41/2018). MJ thank Prof. Manu Agarwal from University of Delhi and Prof. Madan Pal Singh from ICAR-Indian Agricultural Research Institute, New Delhi for their suggestions and infrastructural support in growing plants. All authors thank BITS Pilani for providing the infrastructure and administrative support.

## Author contribution

HS conducted the experiments, analyzed the data and wrote the paper. RS helped in detailed data analysis. MJ conceptualized the idea, received the grant, monitored the project, helped in experiments and wrote the paper. All authors checked and approved the paper.

